# Omega-3 Fatty Acid Derived Neuroactive Lipids - Docosahexaenoyl-Glycine and Its Epoxide Metabolites are Multifunctional Lipid Mediators

**DOI:** 10.1101/2025.10.17.681057

**Authors:** Justin S. Kim, Luca Franchini, Yevgen Yudin, Anna N. Denissiouk, Tibor Rohacs, Cesare Orlandi, Aditi Das

## Abstract

Lipid mediators derived from ω-3 and ω-6 polyunsaturated fatty acids (PUFAs) support neurological health in part through their oxidative and non-oxidative transformation into a diverse array of bioactive molecules. Among these are lipidated neurotransmitters, formed via conjugation of neurotransmitters with fatty acids such as arachidonic acid (AA) or docosahexaenoic acid (DHA). Previous studies links these lipidated neurotransmitters to beneficial outcomes in neurological diseases. Here, we focus on two such endogenous lipidated neurotransmitters, arachidonoyl glycine (NA-Gly) and docosahexaenoyl glycine (DHA-Gly) and demonstrate their further biotransformation by cytochrome P450 enzymes into epoxidized metabolites. These metabolites are structurally multifunctional, combining both epoxide and glycine moieties. In lipopolysaccharide-stimulated microglial cells, we observe increased formation of NA-Gly and DHA-Gly, correlating with their anti-inflammatory effects. Functionally, these lipidated glycines are selective and act as inverse agonists of G protein–coupled receptor 55 (GPR55) and selectively potentiate transient receptor potential vanilloid 4 (TRPV4), but not TRPV1 or TRPM3 channels. Together, our findings identify NA-Gly, DHA-Gly, and their epoxide derivatives as multifunctional lipid mediators with anti-inflammatory properties and selective receptor modulation, positioning them as potential therapeutic leads in neuroinflammation and reinforce the critical side role of glycine in brain function.

**Significance:** Lipidated neurotransmitters derived from omega-3 and omega-6 polyunsaturated fatty acids (PUFAs) contribute to neurological health through their conversion into a diverse array of bioactive signaling molecules. In this study, we study docosahexaenoyl glycine (DHA-Gly) and demonstrate their further enzymatic transformation by cytochrome P450 epoxygenases into epoxidized derivatives. These structurally distinct metabolites exhibit anti-inflammatory activity in microglial cells and interact with GPR55 and TRPV4, but not TRPV1 or TRPM3. Our findings highlight a new class of multifunctional lipid mediators with therapeutic potential for targeting neuroinflammation and related neurological disorders.

## INTRODUCTION

Diet rich in ω-3 fatty acids such as docosahexaenoic acid (DHA) is known to promote neurological health [1]. Neural membranes are rich in DHA. Phospholipids in the gray matter of cerebral cortex and photoreceptor cells in the retina contain DHA [2]. DHA was previously thought to be mostly involved in regulation of physicochemical properties of the neural membrane such as membrane fluidity, permeability and viscosity in synaptic membranes. Recently, it has been postulated that DHA based phospholipids serve as reservoirs to generate DHA based lipidated neurotransmitters such as docosahexaenoyl ethanolamide (DHEA or DHA-EA) (synaptamide), DHA-Dopamine (DHA-DA), DHA-Serotonin (DHA-5HT) [3]. These DHA based neurotransmitters activate signaling pathways that sustain synaptic function, neuronal survival and are neuromodulators [4–6]. The biochemical mechanisms facilitating these effects by DHA-NTs are yet to be fully elucidated. Previously, we reported that DHA is non-oxidatively converted to DHEA and is further metabolized by cytochrome P450s in the brain to form DHEA-epoxides. These DHEA-epoxides, such as 19,20-epoxydocosahexaenoyl-ethanolamide (19,20-EDP-EA) interact favorably with cannabinoid receptor-1 (CB1) and −2 (CB2) [3, 7]. CB1 is predominantly expressed in the central nervous system (CNS), and CB2 is found in both peripheral and CNS immune cells [8–10]. In addition, they reduce inflammatory responses and promote angiogenesis and vasodilation.

Herein we focus our studies on arachidonic acid (AA) and DHA conjugated to glycine in the brain to form N-arachidonyl-glycine (NA-Gly) and docosahexaenoyl-glycine (DHA-Gly) (Figure 1). Glycine by itself is a fundamental building block of proteins and provides nitrogenous backbones to form neurotransmitters, nucleotide bases and hormones [11, 12]. Glycine is found in large quantities in the brain and acts as a primary neurotransmitter of inhibitory interneurons in the spinal cord. Glycine interacts with glycine receptors (GlyR) and is involved in synaptic transmission, motor control, and pain perception [13, 14]. In addition, glycine possesses antioxidant, anti-inflammatory, and immunomodulatory functions [11, 12, 15]. The significance of glycine is highlighted in that low glycine levels suppress immune responses, stunt growth, and cause abnormal metabolism. Glycine is also an agonist of N-methyl-D-aspartate (NMDA) receptors in the brain and crosses the blood-brain barrier following high-dose oral administration. Glycine is also used as an adjuvant therapy to treat patients with neurological diseases such as schizophrenia [16].

**Figure 1.**
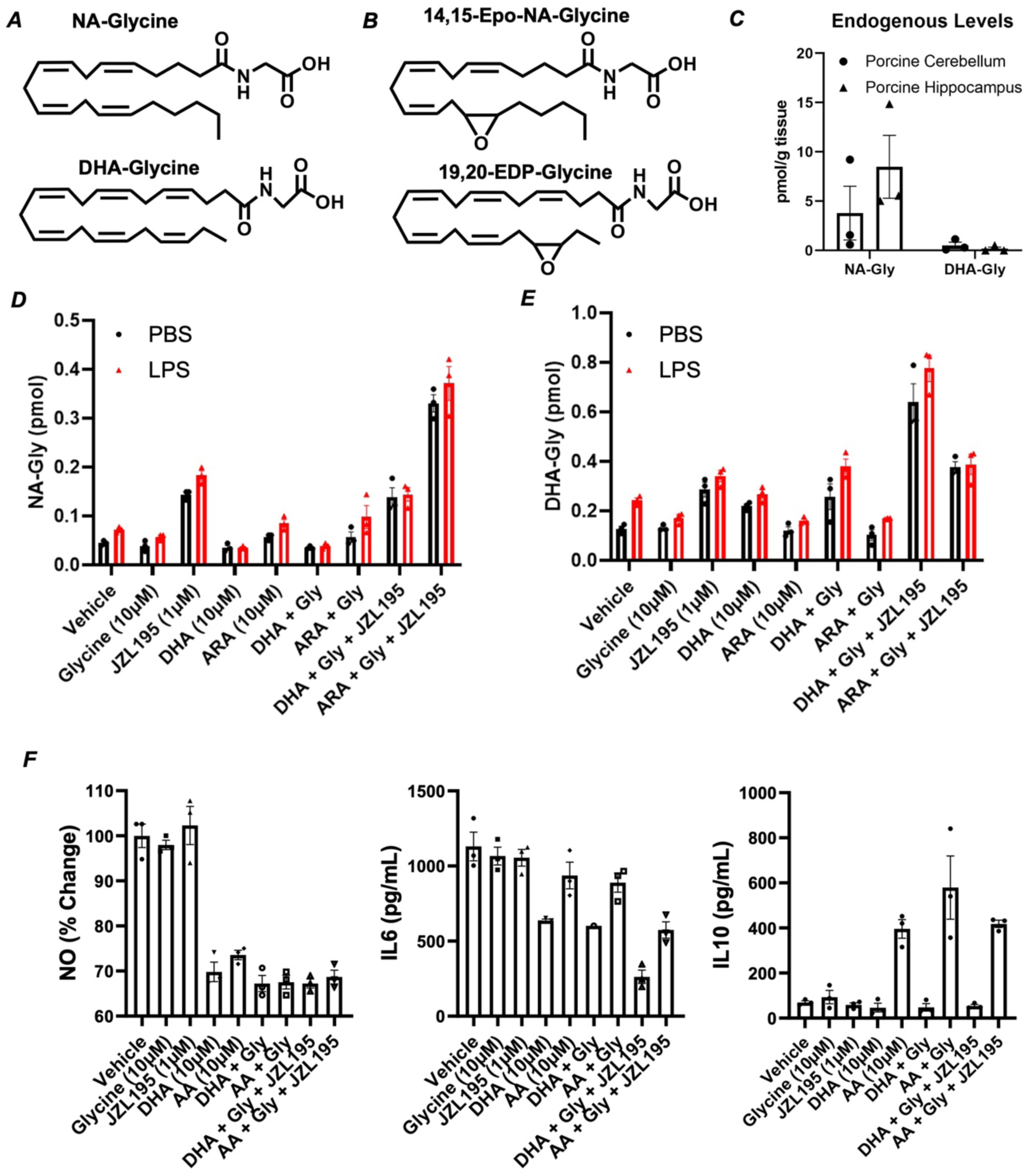
Endogenous levels and production of lipidated glycine molecules. (A) Structures of NA-Glycine and DHA-Glycine, and (B) epoxide metabolites - 14,15-Epo-NA-Glycine and 19,20-EDP-Glycine. (C) Endogenous levels of lipidated glycine in porcine brain regions. (D-E) Formation of NA-Gly and DHA-Gly in BV-2 microglia after feeding with precursor molecules with and without LPS. (F) Measurement of change in Nitric oxide (NO), IL-6 and IL-10 in microglial cells after being treated with the precursor molecules in the presence and absence of LPS. All cell culture experiments were done n=3-5 and errors are reported.

Glycine is lipidoylated to form *N*-acyl amino acids or lipo-amino acids. An example of a *N*-amino acid conjugate is *N*-arachidonoyl-glycine (NA-Gly) by which glycine is conjugated to arachidonic acid via an amide linkage (Figure 1A). Several biosynthetic pathways have been demonstrated for the biosynthesis and degradation of *N*-acyl amino acid conjugates [5, 17–19]. N-acyl glycine is formed from the condensation of the acyl moiety, free fatty acid, or coenzyme A derivative with glycine [20][21]. Besides NA-Gly, there are several structurally similar lipidated amino acids that have been reported before [20]. Overall, the predominant synthesis route responsible for the formation of these lipidated amino acids is open to further study.

Another lipoylated glycine is DHA-Glycine (Figure 1A) that was discovered in the brain, and has been reported to increase in the brain following carrageenan inflammatory stimuli in mice [22]. In addition, N-acyl phosphatidylethanolamine phospholipase D (NAPE-PLD) knockout in *Luquent* lines showed an increase of DHA-Gly in the striatum and cortex in the brain [23]. NAPE-PLD catalyzes the conversion of N-acyl-phosphatidylethanolamine (NAPE) into bioactive signaling molecules like anandamide and oleoyl ethanolamine. DHA-Gly was reported to activate IKS channels, suggesting it may be used to treat hereditary congenital long QT syndrome in the heart [24]. Other studies have used a mixture of *N*-acyl amides which contained DHA-Gly to activate the heat and capsaicin-sensitive Transient Receptor Potential V1 (TRPV1) ion channels, and to a smaller extent TRPV4, an osmo-and heat-sensitive ion channel [22].

In an oxidative metabolism pathway, omega-3 and omega-6 fatty acids are metabolized by eicosanoid-synthesizing enzymes from the cyclooxygenase (COX), lipoxygenase (LOX), and cytochrome P450 epoxygenase (CYP) pathways to generate oxidized lipid metabolites. The acyl chain of NA-Gly and DHA-Gly are site of metabolism by these enzymes to form oxidized molecules that are bioactive. For instance, DHEA was metabolized by LOX and CYP pathway enzymes to form oxidized metabolites that all demonstrated anti-inflammatory properties [3]. Here, we report the metabolism of NA-Gly and DHA-Gly by CYP pathway enzymes to generate novel epoxides that are multifunctional bioactive molecules (Figure 1B).

CYP450s are linked to the regulation of inflammatory diseases through the production of anti-inflammatory lipid-epoxides such as epoxyeicosatrienoic acids [25], epoxyeicosatetraenoyl-ethanolamides and epoxydocosapentaenoyl-ethanolamides [26]. These epoxidated lipids have been demonstrated to be crucial as anti-inflammatory and pro-resolving lipid mediators [26, 27]. The significance of this can be easily highlighted by examining CYP2J2, an epoxygenase that is well-established to be important for cardiac health. For instance, overexpression of CYP2J2 in rodents protected them from doxorubicin-induced cardiotoxicity [28]. More mechanistic understandings of CYP2J2 and anthracyclines were later reported [29–31]. Collectively, these studies demonstrate that arachidonic acid must be released and metabolized by CYP450s to elicit its effects on our body.

NA-Gly and DHA-Gly have chemical structure like endocannabinoids such as anandamide (AEA) and docosahexaenoyl ethanolamine (DHEA). While AEA and DHEA activate cannabinoid receptors and are anti-inflammatory in their own capacity, they are further metabolized to downstream lipid metabolites as explained before. Therefore, it is important to show whether NA-Gly and DHA-Gly are metabolized by cytochrome P450s to form epoxide metabolites (Figure 1B). AD-STOP

CYP450s have varying levels of expression over time following an inflammatory stimuli in RAW264.7 macrophages [32]. Recently, CYP4F2 was reported to be a human-specific determinant of circulating *N*-acyl amino acid levels in the body [33]. Although NA-Gly is not uniformly distributed in the body [34, 35], there is expression in regions where the regulation of inflammatory process is critical. For example, NA-Gly and DHA-Gly are both detected in the central nervous system. Previous study demonstrated that NA-Gly and DHA-Gly both increased in the striatum, hippocampus, and cerebellum 3-hours post carrageenan stimulation in rodents [22]. These suggest that the lipidated glycines may be released to elicit an anti-inflammatory or pro-resolving effect following inflammatory stimuli. Separately, the levels of the lipidated glycines were elevated in NAPE-PLD KO mice, compared to other lipidated neurotransmitters such as N-acyl ethanolamine [23]. Considering the general dysregulation of CYP450s and the increase of lipidated neurotransmitters in disease states, it is plausible there is lipid release and subsequent metabolism by CYP450s to form bioactive pro-resolving lipid mediators.

NA-Gly has been shown to suppress pain responses. For instance, NA-Gly reduced thermal hyperalgesia [36] and nerve injury-induced neuropathic pain [37] through intrathecal injection in inflammatory pain models. The analgesic effects of NA-Gly in rodent models were not inhibited by cannabinoid receptor antagonists, suggesting alternative receptors. NA-Gly and alternative receptor interactions have been reported with mixed findings. NA-Gly does not activate cannabinoid receptors but exhibits anti-inflammatory and anti-nociceptive properties through other receptors [38–40]. NA-Gly was identified as an endogenous ligand for G-Protein Coupled Receptor 18 (GPR18) [41] which was termed as the NA-Gly Receptor. The hypothesis that NA-Gly and similar *N*-acyl amides are ligands of orphan GPCRs has been explored previously [4]. However, there is uncertainty within the field regarding NA-Gly receptor interactions. For instance, while NA-Gly was first reported to be an endogenous ligand for GPR18, several other studies report that NA-Gly does not act as an agonist of GPR18 [42, 43]. Recently, NA-Gly was suggested as an endogenous ligand for orphan GPR55 [44]. Another receptor that NA-Gly or DHA-Gly can interact with is glycine transporter type 2 (GlyT2) which regulates the transport and concentration of glycine entering the synaptic cleft [45–47]. As GlyT2 has been linked to chronic pain, lipid-based inhibitors have become a research target as a potential treatment [48]. While the mechanism is not clear, NA-Gly inhibited formalin-induced pain responses by inhibiting GlyT2 [34]. Non-canonical GPCRs such as GPR18 were also investigated due to their ability to heteromerize with cannabinoid receptors. It was discovered that NA-Gly stimulates orphan GPR18, GPR92, and GPR55 [49]. As GPR92 co-localizes with transient receptor potentiated vanilloid receptor 1 (TRPV1), a channel that is associated with reducing pain responses, we hypothesize that lipidated-glycine is a TRPV agonist.

With regards to formation and degradation of lipid-amino acid conjugates, several routes have been proposed. One such enzymatic route is via fatty acid amide hydrolase (FAAH) that can hydrolyze endocannabinoids and amidated lipid conjugates [50]. FAAH is known to hydrolyze anandamide [51], an endocannabinoid lipid mediator that acts as an endogenous ligand of CB1 receptors. As endocannabinoids mediate pain, and neurodegenerative diseases, there is high interest to develop FAAH inhibitors. While there are synthetic inhibitors of FAAH to increase endogenous levels of fatty acid ethanolamides [52], naturally occurring substrates or inhibitors of FAAH contribute to the higher abundance of endocannabinoids. Interestingly, FAAH also has wide substrate specificity. For instance, in rat ventrolateral periaqueductal gray, FAAH inhibition increased 2-arachidonoyl glycerol [53]. In a separate study administered FAAH inhibitor cyclohexylcarbamic acid 3-carbamoyl biphenyl-3-yl ester (URB597) in rats and discovered an increase of NAEs anandamide and oleoyl ethanolamide, as well as 2-arachidonoyl glycerol [54]. These findings show that FAAH traditionally thought to hydrolyze anandamide and related molecules also hydrolyzed 2-AG which does not contain an amide linkage [19, 55]. As glycine is abundant in the central nervous system, other groups have examined the analgesic effects of NA-Gly in neuropathic pain models. For instance, NA-Gly reduced thermal hyperalgesia [36] and nerve injury-induced neuropathic pain through intrathecal injection in inflammatory pain models. Surprisingly, it was reported that analgesic effects of NA-Gly in rodent models were not inhibited by cannabinoid receptor antagonists, suggesting interactions with other receptors. Through the process of competitive inhibition, many substrates of FAAH can act as inhibitors of endocannabinoid hydrolysis. Therefore, we evaluate if NA-Gly and DHA-Gly are natural endogenous inhibitors of FAAH.

Another enzyme, Peptidase M20 Domain containing 1 (PM20D1) can also regulate the levels of bioactive N-acyl amide lipids. PM20D1 is one of five enzymes in the 20D family of peptidases that can act as both hydrolase and synthase of lipidated amino acids [56]. The hydrolase and synthase activities were observed in C18:1 lipid conjugated to various amino acids including phenylalanine, leucine, valine, isoleucine, methionine, serine, glycine, and many more [56]. A more recent study further identified that FAAH and PM20D1 may cooperatively regulate the levels of N-acyl amino acids [19]. This study observed hydrolase and synthase activity in lipids conjugated to similar amino acids. Importantly, these studies suggest a bidirectional role of FAAH and PM20D1 by which they can act as both hydrolase and synthase. While these enzymes are commonly thought to be involved strictly in the degradation process of lipidated amino acids and endocannabinoids, the knowledge of how they can act as a synthase is equally important.

As there is an abundance of glycine, AA and DHA in the central nervous system, we investigate lipidated neurotransmitters N-arachidonoyl-glycine (NA-Gly) and docosahexaenoyl-glycine (DHA-Gly) with respect to their formation, receptor interactions and further metabolism. They are naturally found in the brain. They are produced by unknown pathway and potentially by FAAH and PM20D1 and interact through receptors such as GPR18, GPR55. Furthermore, they are also target for oxidative enzymes such as cytochrome P450s that are upregulated during inflammation to form more potent anti-inflammatory DHA-Gly epoxides (19,20-EDP-Gly) and NA-Gly epoxides (14,15-EET-Gly) (Figure 1B). Overall, we report that arachidonic acid (AA) and DHA are conjugated to glycine in the brain to form N-arachidonyl-glycine (NA-Gly) and docosahexaenoyl-glycine (DHA-Gly), which is subsequently converted by cytochrome P450s to form epoxides in a neuroinflammatory state.

## RESULTS

### Endogenous presence of NA-Gly and DHA-Gly in porcine brain

We first measured the endogenous presence of lipidated amino acids and neurotransmitters. For this, we generated a quantitative LC-MS/MS panel using authentic standards. Two extraction solvent conditions were utilized to ensure maximal extraction of lipidated glycine (see methods). Using ethyl acetate: hexane, NA-Gly and DHA-Gly are present in the cerebellum at 4 pmol/g tissue and 1 pmol/g tissue, respectively (Figure 1C). In the hippocampus, NA-Gly is present at 8 pmol/g tissue, while DHA-Gly is below detection limits (Figure 1C). The trend is that NA-Gly is detected at higher concentrations than that of DHA-Gly in both the cerebellum and hippocampus. In addition, the concentration of the lipidated-glycine is higher in the hippocampus than cerebellum in ethyl acetate: hexane-extracted tissues. Similar patterns were observed when using extraction solvent chloroform: methanol. NA-Gly was only detected in the hippocampus at 3 pmol/g tissue. NA-Gly was not detected in the cerebellum, and DHA-Gly was not detected in either cerebellum or hippocampus. These findings demonstrate that these molecules are endogenously found in tissues, but their levels are variable and is dependent on extraction methodology and columns used in the process, the time of tissue extraction and rate of molecule turnover. It is important to note the presence of these molecules endogenously and localization to specific tissues warrants further inquiry into their formation and metabolism.

### Microglia synthesize NA-Gly and DHA-Gly under inflammatory conditions

Microglia are the brain’s resident immune cells and once they are activated under inflammatory conditions, they produce various lipid mediators that can either promote or resolve inflammation. Changes in microglial lipid metabolism are linked to shifts in their function from protective to harmful and contribute to neurodegenerative diseases. Here we studied whether BV-2 microglia can synthesize NA-Gly or DHA-Gly under non-inflammatory and inflammatory conditions in the presence and absence of supplementation with the precursors – glycine and lipids. We used JZL195 as a potent inhibitor of fatty acid amide hydrolase (FAAH), an enzyme that can hydrolyze amide bonds located at the fatty acid head group moiety. We treated the cells with glycine, JZL195, DHA, arachidonic acid (ARA), DHA + Gly, ARA + Gly, DHA + Gly + JZL195, or ARA + Gly + JZL195. We observe that NA-Gly and DHA-Gly levels increase with the supplementation of their precursor molecules. We see an increase upon feeding with the fatty acid precursor ARA or DHA (Figure 1D-E). This is evident by the fact that ARA or DHA supplementation increases NA-Gly and DHA-Gly. Furthermore, the quantity of NA-Gly and DHA-Gly increasing in the presence of JZL195 is likely due to the inhibition of FAAH hydrolase activity of the lipidated-glycine molecules that we study later.

### Signaling Interactions of Docosahexaenoyl-Glycine and N-Arachidonoyl-Glycine with Cannabinoid Receptors

Using a sensitive Bioluminescence Resonance Energy Transfer (BRET)-based (G protein nanoBRET) assay we obtained a fingerprint of the G protein coupling profile of cannabinoid receptors (CB1R, CB2R, GPR55, and GPR119) in response to agonist stimulation [57]. In the G protein nanoBRET assay, HEK293 cells are transiently transfected with 5 components including the GPCR of interest, a wild-type Gα protein, Gβ1 and Gγ2 subunits fused to a split fluorescent protein (Venus, acceptor), and the C-terminus portion of the Gβγ effector GRK3 fused to a luciferase (GRK3-CT-nanoluc, donor). Stimulation of a GPCR results in the dissociation of heterotrimeric G proteins allowing Gβγ-Venus to interact with GRK3-CT-nanoluc and produce a BRET signal that is an index of receptor activation. Here, we first confirmed the activation of CB1R, CB2R, GPR55, and GPR119 when co-expressing G proteins that are reported to be activated by each receptor applying established synthetic agonists (Figure 2A). Then, we obtained concentration-response curves exploring the activity of DHA-Gly on the same four receptors (Figure 2B). None of the receptors showed activation in response to DHA-Gly application. We next verified that other G proteins were not activated by the application of a saturating concentration of DHA-Gly (Figure 2C). Similarly, we explored the effect of NA-Gly on the activation of the CB1R, CB2R, GPR55, and GPR119. We found that cognate G proteins were not activated at any concentration applied (Figure 2D). The effect on the activation of non-cognate G proteins was also undetectable by our G protein nanoBRET assay (Figure 2E). Overall, this set of data suggests that DHA-Gly and NA-Gly do not function as cannabinoid receptor agonists.

**Figure 2.**
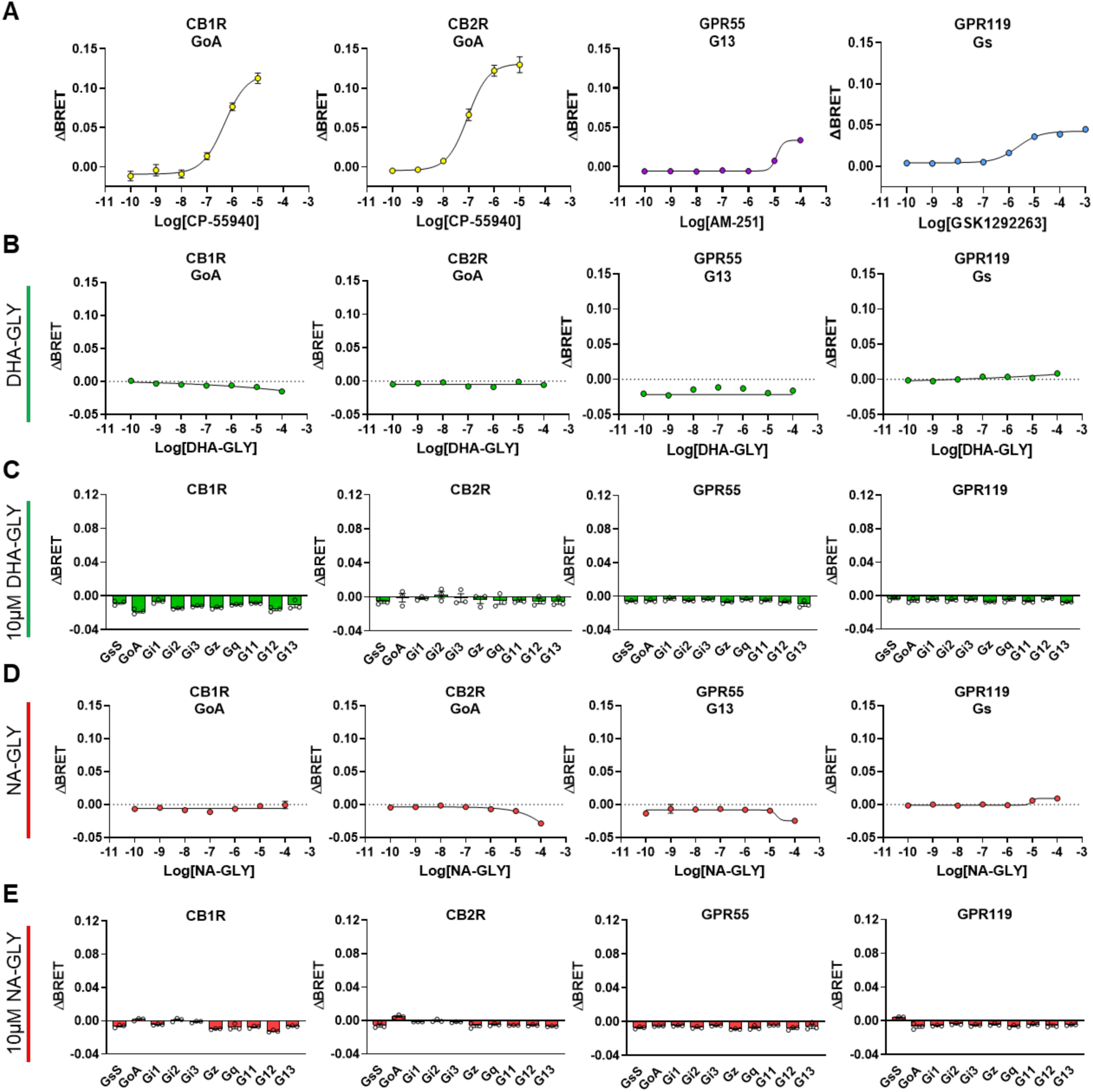
Activation of cannabinoid receptors. **(A)** Concentration-response curves for CB1R, CB2R, GPR55, and GPR119 obtained using G protein NanoBRET assay with respectively coupled Gα proteins (GoA for CB1R and CB2R; G13 for GPR55; Gs for GPR119). The following synthetic agonists were applied: CP-55940 CB1R and CB2R; AM-251 for GPR55, and GSK-1292263 for GPR119. **(B)** Concentration-response curves for indicated cannabinoid receptors in response to DHA-Gly application. **(C)** Individual G protein activation in response to 10µM DHA-Gly. **(D)** Concentration-response curves for indicated cannabinoid receptors in response to NA-Gly application. **(E)** Individual G protein activation in response to 10µM NA-Gly. N=3.

### NA-Gly is an inverse agonist of GPR55

We next sought to investigate the possible role of DHA-Gly and NA-Gly as inverse agonists of cannabinoid receptors (CB1R, CB2R, GPR55, and GPR119). To this goal, we transfected cells with each of the four cannabinoid receptors and the respective BRET sensors. Cells were pre-treated with indicated concentrations of respective agonists and then vehicle, 50µM NA-Gly, or 50µM DHA-Gly were applied (Figure 3). We observed that treatments with NA-Gly significantly inhibited the BRET signal generated by activation of the four cannabinoid receptors, with GPR55 showing the greatest effect (Figure 3C). DHA-Gly showed comparable effects to vehicle treatments (Figure 3). As an additional control of specificity, we treated a non-cannabinoid receptor, α2-adrenergic receptor (ADRA2A) (Figure 3E). The size of the effect of 50µM NA-Gly on ADRA2A (-34%) was significantly lower than what was observed for GPR55. Finally, we used an orthogonal assay to establish the effect of NA-Gly as an inverse agonist for GPR55. In this case, we used an SRE luciferase reporter where luciferase accumulation in response to G12/13 receptor activation is measured. As expected, overexpression of constitutively active GPR55 significantly induced luciferase accumulation, and treatments with the 100µM of GPR55 agonist AM-251100 further promoted luciferase accumulation (Figure 3F). Importantly, treatments with NA-Gly completely abolished luciferase accumulation induced by GPR55 constitutive activity or agonist-induced activity (Figure 3F). Altogether, these results strongly support an effect of NA-Gly as an effective inverse agonist of GPR55.

**Figure 3.**
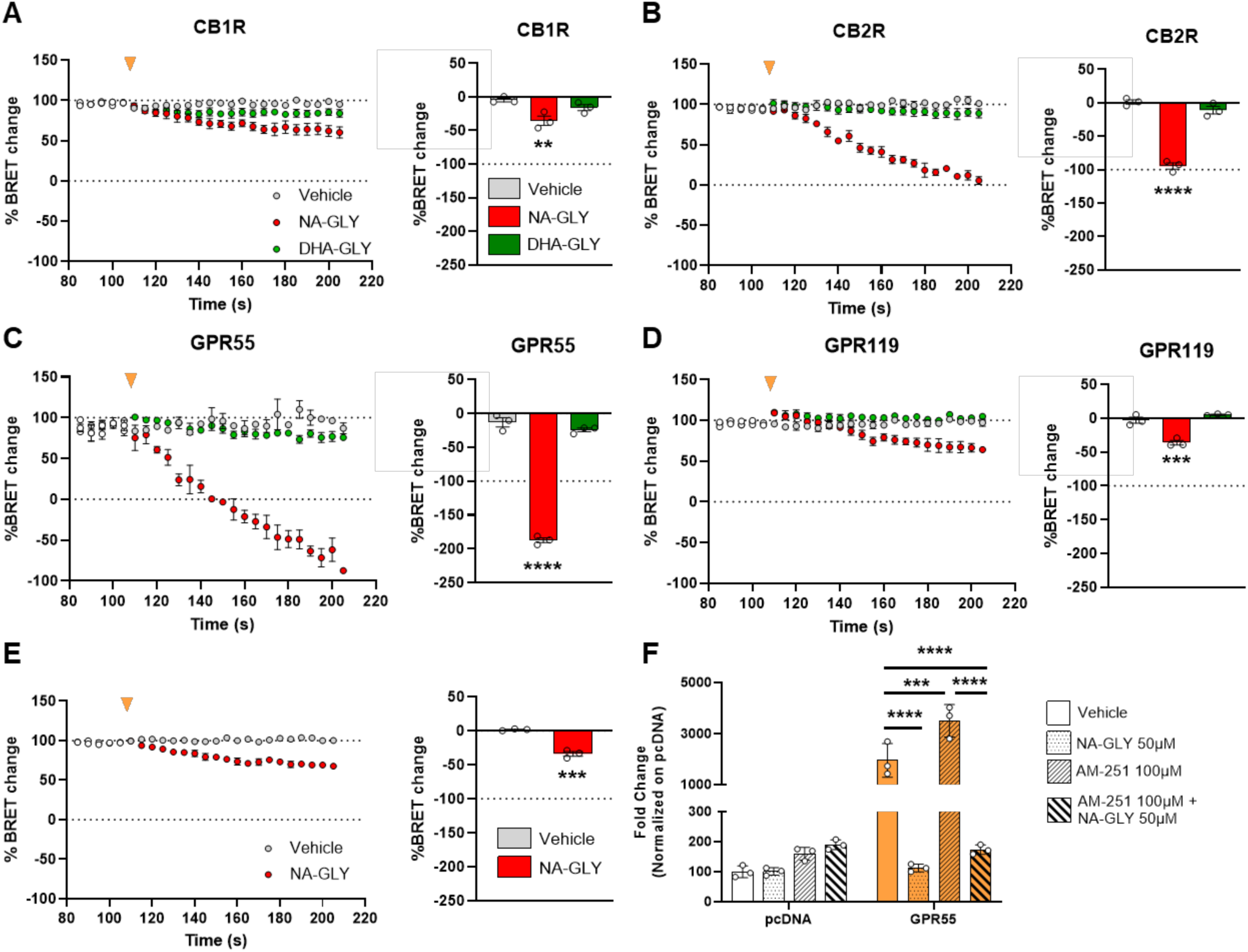
Inhibition of cannabinoid receptors by NA-Gly and DHA-Gly. HEK293T cells were transfected with NanoBRET sensor, indicated receptors, and each coupled Gα protein: GoA for CB1R **(A)**, GoA for CB2R **(B)**, G13 for GPR55 **(C)**, and Gs for GPR119 **(D)**. Cells were treated with the following agonists: 1µM CP-55940 for CB1R and CB2R; 100µM AM-251 for GPR55; 1μM GSK-1292263 for GPR119. After reaching plateau values vehicle, 50µM NA-Gly, or 50µM DHA-GLY were applied (yellow arrowhead) and ΔBRET was recorded for 100s. Bar graphs report the percentage of change in ΔBRET after vehicle, NA-Gly, or DHA-Gly application for each receptor. N=3, One-way ANOVA, Dunnett’s multiple comparison vs vehicle. **p<0,01; ****p<0.0001. **(E) Inhibition of α2A adrenergic receptor (ADRA2A) by NA-Gly**. Cells transfected with Nano-BRET sensors, GoA, and ADRA2A were stimulated with 0.1μM clonidine. After reaching maximal values, vehicle or 50μM (final concentration) NA-Gly was applied (yellow arrowhead) and ΔBRET was recorded for 100s. The bar graph (right) reports the quantification of the percentage of change in ΔBRET after treatment. N=3; unpaired t-test; *** p<0.001.

### NA-Gly and DHA-Gly Inhibits Fatty Acid Amide Hydrolase

Fatty acid amide hydrolase is an enzyme that is known to degrade other endocannabinoids such as anandamide by acting on the amide bond. In Figure 1, our results demonstrate that supplementation with the fatty acid precursors ARA and DHA enhances the cellular production of NA-Gly and DHA-Gly, respectively. This increase is further amplified in the presence of the FAAH inhibitor JZL195, suggesting that FAAH contributes to the hydrolytic breakdown of these lipid-glycine conjugates. Together, these findings highlight the importance of precursor availability and FAAH activity in regulating endogenous levels of NA-Gly and DHA-Gly by potentially acting as a substrate of FAAH as they both contain an amide bond. We conducted a FAAH inhibition assay with DHA-Gly and NA-Gly in a dose-dependent manner (1μM – 100 μM) to determine if they can act as inhibitors of FAAH-mediated AMC-Anandamide degradation. We determined that both NA-Gly and DHA-Gly inhibit FAAH in a dose-dependent manner. In addition, NA-Gly has greater affinity and inhibition potential than DHA-Gly which can be observed by the higher percent inhibition across all concentrations (Figure 4). Following a 4-parametric logistic regression fitting, we confirmed these observations and determined that the IC50 of NA-Gly and DHA-Gly was 14.20 ± 1.07 μM and 35.35 ± 1.82 μM, respectively.

**Figure 4.**
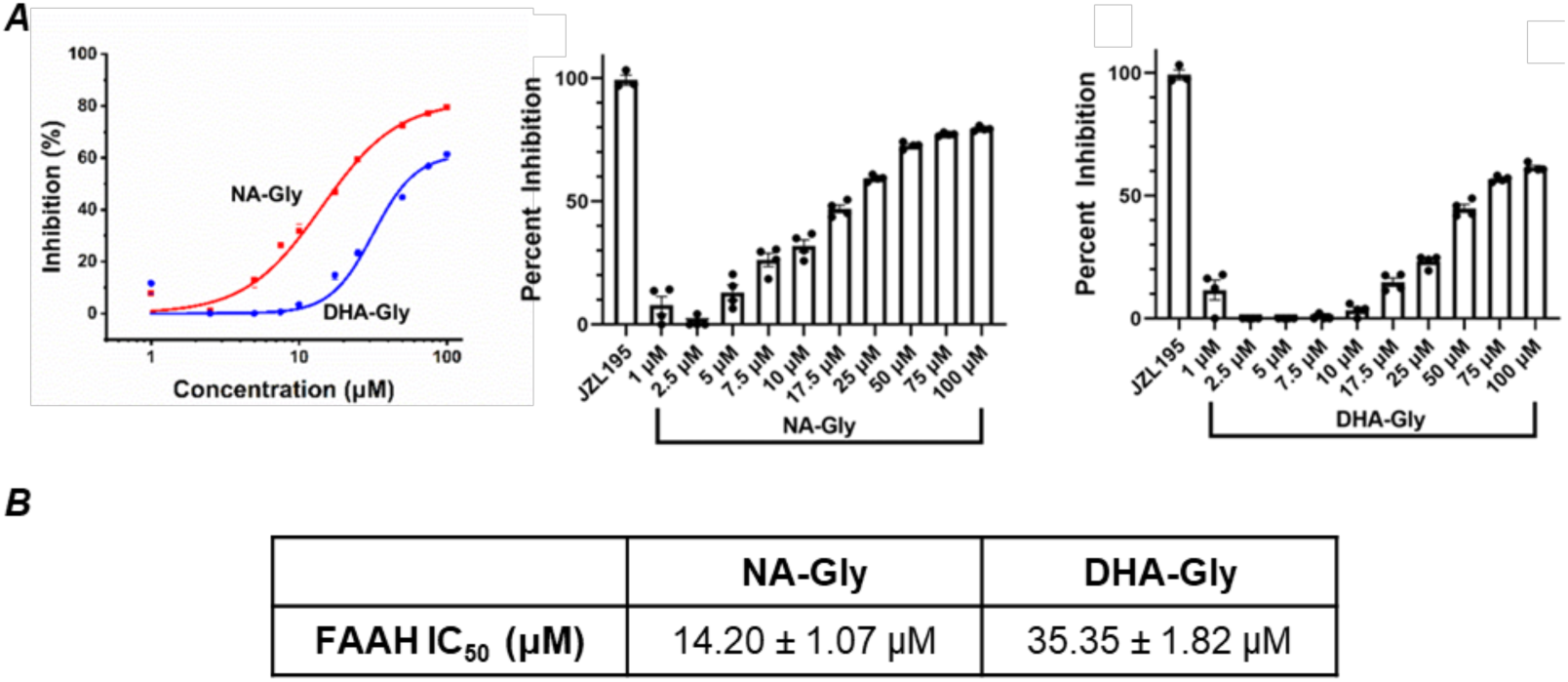
Fatty Acid Amide Hydrolase Inhibition. NA-Gly and DHA-Gly potential to inhibit Fatty Acid Amide Hydrolase (FAAH) measured by release of fluorophore from anandamide-AMC. (A) Concentration dependent increase of FAAH inhibition by NA-Gly and DHA-Gly. (B) Summary of IC50 (μM) values based on response curve. N=3 per concentration.

### NA-Gly and DHA-Gly are substrates for metabolism by cytochrome P450 epoxygenases

***CYP450 expression increases in BV-2 microglia following inflammatory stimuli.*** Upon stimulation by LPS, BV-2 microglia generate an inflammatory response in a time-dependent response. We also investigated the time dependent expression of CYP450 epoxygenases in microglia. Interestingly, CYP2J6, a rodent CYP450 epoxygenase with greatest homology similarity to human CYP2J2 epoxygenase, had several fold increase at 18 and 24 hours following LPS-stimulation (Figure S1). These data highlight the potential role of the increased CYP2J6 expression to the production of epoxide metabolites.

***NA-Gly and DHA-Gly binds to CYP450 epoxygenases.*** As epoxide formation is dependent on epoxygenase-mediated metabolism, we investigated whether they could bind to CYP450s that are found in the central nervous system and possess epoxygenase activity. We showed that NA-Gly and DHA-Gly bind to human CYP2J2 tightly (Figure S2). The lipidated glycines exhibited a Type I substrate binding to CYP2J2, and a Type II substrate binding to CYP3A4. With regards to CYP2J2, NA-Gly and DHA-Gly bind with KD = 3.30 ± 0.31 μM, and KD = 2.47 ± 0.26 μM, respectively. The lipidated glycines bind to CYP3A4 weaker than they bind to CYP2J2. Specifically, NA-Gly and DHA-Gly bind to CYP3A4 with KD = 30.04 ± 16.65 μM and KD = 20.31 ± 6.13 μM (Figure S2). CYP3A4 is primarily a xenobiotic metabolizing CYP450 and is not known for binding lipids. Therefore, it is not surprising that the lipidated glycines bind to CYP3A4 with weaker affinity.

***Metabolism of DHEA, NA-Gly, and DHA-Gly indicate epoxide products.*** While binding indicates there is ligand-receptor interactions, we performed a metabolism analysis to confirm whether CYP450s metabolize substrates to form epoxides (Figure 5A and 5B). Metabolism of DHA-glycine by rat brain microsomes (Figure 5C and 5D), human liver microsomes (Figure S7), and recombinant CYP2J2 (Figure 5), CYP3A4 (Figure S5), and CYP2D6 (Figure S6) were analyzed on full-scan LC-MSMS. Following mass spectrometry analysis, there were corresponding masses to the parent molecule of 372.28 originating from the 6.23 minute peak on the LC chromatogram. Importantly, there were corresponding DHEA-Epoxide *m/z* values of M + H^+^ = 388.31, which correlates with previously reported findings. NA-Gly and DHA-Gly have masses of 362.26 and 385.26, respectively. Following metabolism of NA-Gly and DHA-Gly by rat brain microsomes, human liver microsomes, and recombinant CYP450s, the parent masses were found on the MS chromatogram with *m/z* corresponding to M + H^+^ = 363.26, and M + H^+^ = 386.26, respectively (Figure 5). Importantly, we first report the product with *m/z* corresponding to M + H^+^ of an epoxide product, similar to that of DHEA. These correlate to the *m/z* of M + H^+^ = 378.26 and M + H^+^ = 402.26 for DHA-Gly and NA-Gly, respectively. These data suggest that NA-Gly and DHA-Gly get epoxidized by epoxygenase enzymes as demonstrated by recombinant CYP450s tested including CYP2J2 (Figure 5A and Figure 5B), CYP3A4 (Figure S5), and CYP2D6 (Figure S6). Importantly, these findings are confirmed by the data from human liver microsomes (Figure S7) and rat brain microsomes (Figure 5C and Figure 5D).

**Figure 5.**
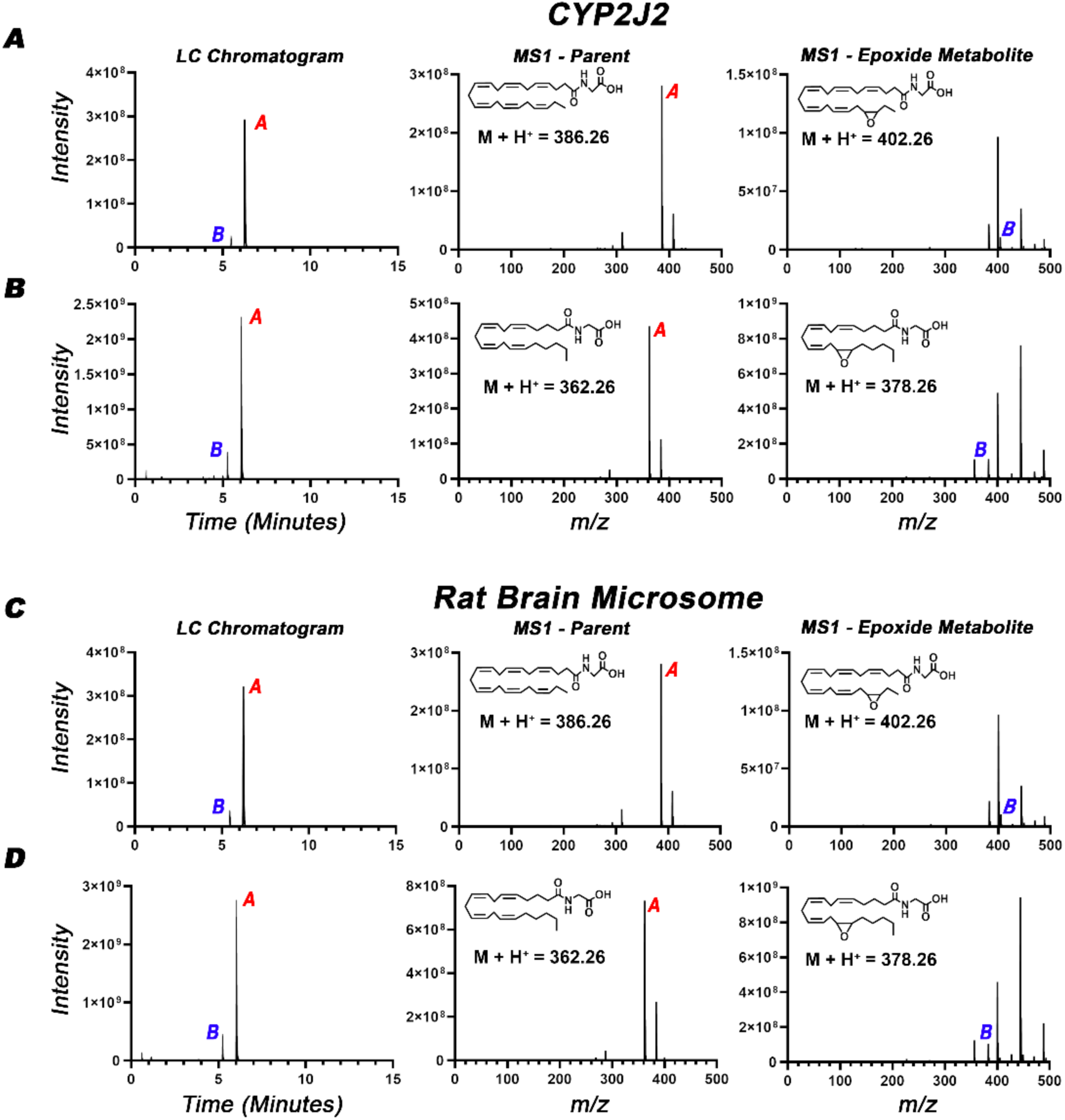
Metabolism of lipidated glycine by CYP450s shows production of epoxide metabolites. CYP2J2-mediated formation of (A) EDP-Glycine from DHA-Glycine, and (B) epo-NA-Glycine from NA-Glycine. Rat brain microsome-mediated formation of (C) EDP-Glycine from DHA-Glycine, and (D) epo-NA-Glycine from NA-Glycine.

***HLM-Metabolism of DHEA, NA-Gly, and DHA-Gly indicate glucuronidated products.*** While epoxidation is a form of Phase 1 substrate metabolism, Phase 2 metabolism can also occur. Phase 2 metabolism is the conjugation of drugs or metabolites to other molecules, and includes processes such as acylation, amino acid conjugation, and glucuronidation. Glucuronidation is a key component of Phase 2 metabolism and requires Uridine 5’diphospho-glucuronosyltransferase (UDP-Glucuronosyltransferase, or UGT). Glucuronidated products are generally more hydrophilic to aid in molecule transport for excretion, and non-drug substrates can be hormones, prostaglandins, or fatty acids. Since glucuronidation requires UGT, we performed fragmentation analysis on the glucuronidated products of our human liver microsome-mediated metabolism of DHEA, NA-Gly, and DHA-Gly. Interestingly, we discovered that there are *m/z* peaks of glucuronidated DHA-Gly, glucuronidated 19,20-EDP-Gly, glucuronidated 19,20-DiHDPA-Gly, glucuronidated 19-HDPA-Gly, and 20-Glucuronidated 19-HDPA-Gly-glucuronate. These peaks correspond to the *m/z* value of M + H^+^. (Table S1 and S2)

### DHA-Glycine and its metabolites are anti-inflammatory in microglial cells

In figure 5, we show that DHA-Gly is metabolized by epoxygenase CYP450s to form epoxy-docosahexaenoyl glycine. To evaluate pharmacology of the epoxy-DHA-Gly, we synthesized and purified 19,20-epoxy-docosahexaenoyl glycine (EDP-Gly). Spectral analysis demonstrates a peak absorbance at 192 nm, which differs from NA-Gly and DHA-Gly which absorb at 194 nm and 195 nm, respectively. Direct injection of purified EDP-Gly onto a triple-quadrupole mass spectrometer shows a mass corresponding to EDP-Gly with *m/z* of M + H^+^ = 402.2. 400Mhz H^+^ NMR was performed with methanol-d4 as the solvent (Figure S3: synthesis scheme and Figure S4 includes characterization)

***Precursors to NA-Gly and DHA-Gly are anti-inflammatory in BV-2 Microglia.*** Since BV-2 microglia fed with the precursors of lipidated-glycine molecules showed an increase of NA-Gly and DHA-Gly, we questioned whether supplementation also decreased inflammation levels when BV-2 microglia are stimulated with LPS. We show that glycine and JZL195 by themself do not decrease nitric oxide production in LPS-stimulated microglia compared to vehicle control (Figure 1F). Interestingly, the supplementation of fatty acids and subsequent combination treatments by which fatty acids are present decreased the levels of nitric oxide production by nearly 30%. Collectively, these data show that JZL195, a fatty acid amide hydrolase inhibitor, is not enough to be anti-inflammatory by itself. Rather, fatty acid supplementation so that cells can synthesize the lipidated glycine and similar molecules is important as demonstrated by the corresponding decreased nitric oxide levels to increased lipidated-glycine levels under fatty acid supplementation.

Cytokine IL6 is a strong indicator of microglial phenotype and its resting or reactive inflammatory state. Precursor feeding highlights a clear trend and corroborates our findings that supplementation with fatty acids is more crucial than glycine supplementation for anti-inflammatory potential in BV-2 microglia. For instance, DHA and DHA + Gly supplementation conditions both reduce IL6 levels, however there is no discernible difference between the two (Figure 1F). Interestingly, and likely due to the increased accumulation of DHA-Gly as FAAH is inhibited by JZL195, cells supplemented with DHA + Gly + JZL195 had the greatest reduction of IL6 levels, from approximately 1100 pg/mL (vehicle) to 250 pg/mL. Interestingly, the same trend is observed with arachidonic acid supplementation conditions however to a lesser degree. Collectively, these data show that fatty acid supplementation to BV-2 microglia produces lipidated glycine molecules that are anti-inflammatory, and DHA supplementation possesses more potent anti-inflammatory potential than that of ARA supplementation. Surprisingly, IL10, an anti-inflammatory cytokine was greatly increased in any feeding condition including arachidonic acid. While it is unclear why IL10 increases with arachidonic and not docosahexaenoic acid supplementation, it is likely there are receptor specificities and various mechanisms of action of both the metabolites produced by BV2 microglia.

***NA-Gly, DHA-Gly, and EDP-Gly Inhibit Microglial IL6.*** Previous studies have illustrated the anti-inflammatory potential of endocannabinoids such as DHEA (DHA-ethanolamine) in LPS-stimulated microglia [26]. Therefore, we assessed the levels of IL6 in LPS-stimulated BV-2 cells. We find that NA-Gly nearly inhibits, or completely inhibits the production of IL6 at 5 and 10 μM treatments (Figure 6A). Similarly, we find that DHA-Gly at 5 and 10 μM treatments nearly inhibits complete production of IL6 in BV-2 microglia stimulated with LPS (Figure 6A). In comparison to other anti-inflammatory molecules, we and others have previously published, lipidated glycine molecules are strong anti-inflammatory molecules on LPS-stimulated microglia. These findings support previous literature suggesting lipidated amino acids such as NA-Gly are anti-inflammatory molecules. We first report EDP-Gly which has not been previously reported. We find that EDP-Gly also nearly inhibits IL6 production in BV-2 microglia at 5 and 10 μM (Figure 6A).

**Figure 6.**
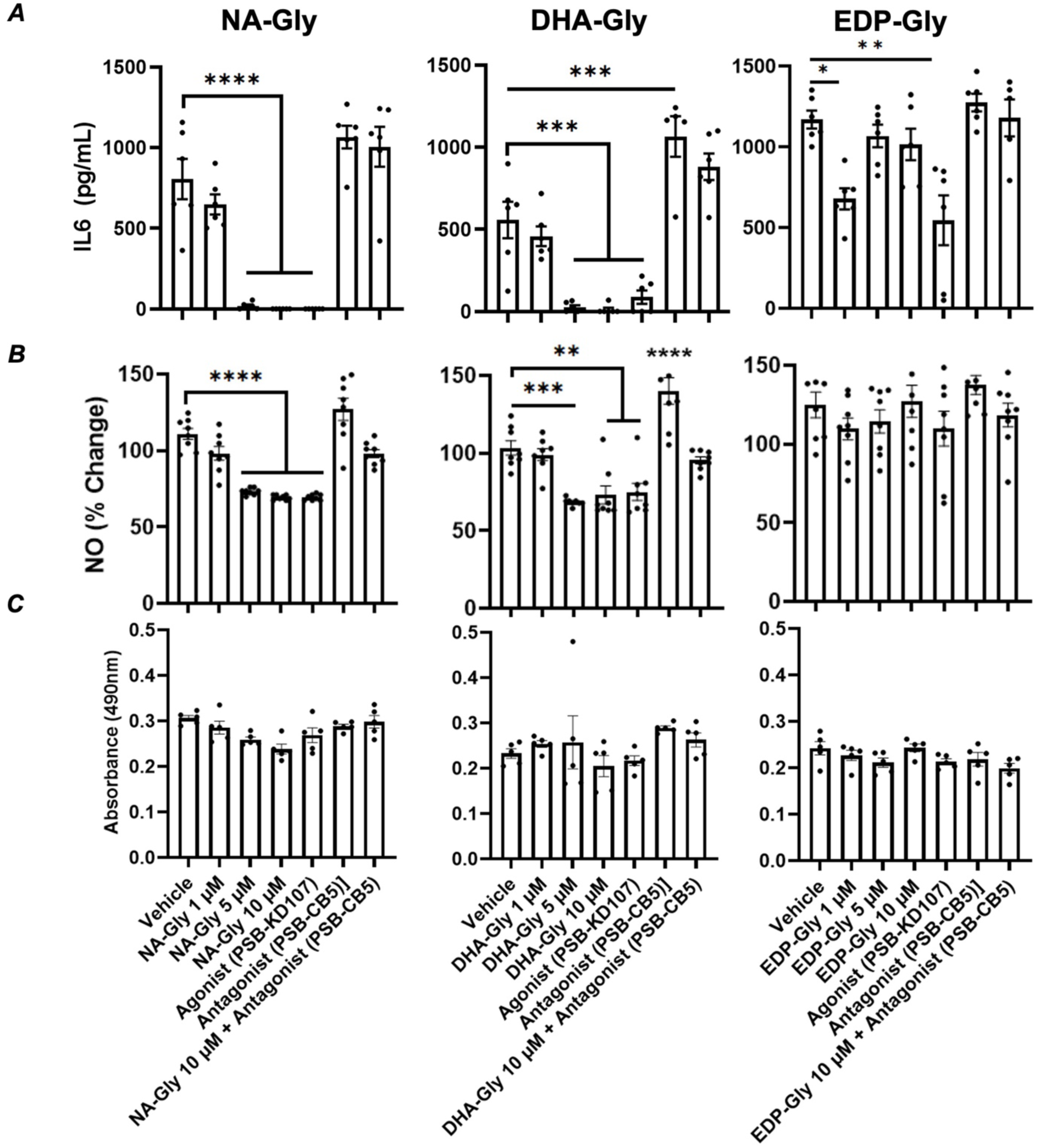
Anti-inflammatory properties of NA-Gly, DHA-Gly and EDP-Gly in LPS-stimulated BV-2 microglia. **(A)** Pro-inflammatory cytokine IL-6 is reduced by NA-Gly, DHA-Gly, and EDP-Gly at 5 ìM and 10 ìM concentrations. GPCR 18 Agonist PSB-KD107 comparably reduces IL6 levels. **(B)** Nitric Oxide is reduced by NA-Gly and DHA-Gly at both 5 μM and 10 μM concentrations. EDP-Gly does not reduce NO levels. **(C)** Lipidated glycines do not induce apoptosis as measured by lactate dehydrogenase. n=5-6 per group. One-way ANOVA statistical analysis was performed.

As there were conflicting data on whether NA-Gly is an agonist of G-protein coupled receptor 18 (GPR18), we conducted LPS-stimulated BV-2 microglial studies with or without the agonist of GPR18. We utilized PSB-KD107 as the agonist, and PSB-CB5 as the antagonist of GPR18. In non-LPS stimulated cells neither PSB-KD107 or PSB-CB5 produced IL6 indicating that these molecules are not pro-inflammatory at 1μM treatments. In LPS-stimulated microglia, we observed nearly a complete inhibition of IL6 by PSB-KD107, and agonist of GPR18. In comparison, we performed dual treatments of NA-Gly and PSB-CB5, DHA-Gly and PSB-CB5, or EDP-Gly and PSB-CB5. When antagonist was present, we observed an increase of IL6 production suggesting that NA-Gly, DHA-Gly and EDP-Gly elicits some of their anti-inflammatory responses through GPR18 receptor in BV-2 microglia.

***NA-Gly, DHA-Gly, and EDP-Gly Inhibit Microglial Nitric Oxide.*** Nitric oxide is an important indicator of inflammatory responses in immune cells including BV-2 microglia. Under LPS-stimulation, NA-Gly, DHA-Gly and EDP-Gly all reduced the percent of nitric oxide present in comparison to the vehicle control (Figure 6B). These correspond with the IL6 levels as reported above. Importantly, GPR18 agonist possessed the same degree of nitric oxide reduction, GPR18 antagonist PSB-CB5 reverted the beneficial effects of the lipidated molecules.

***NA-Gly, DHA-Gly, and EDP-Gly are non-toxic to the cells.*** Considering IL6 and nitric oxide levels were nearly completely ameliorated at 5 and 10 μM concentrations of NA-Gly, DHA-Gly and EDP-Gly, we questioned whether it was due to induced cell death. Importantly, there are no significant differences in the levels of cell death as measured by the lactate dehydrogenase assay (LDH), evident from the non-significant absorbances in all treatment conditions (Figure 6C). Collectively, these data indicate that these lipidated molecules are ant-inflammatory through a non-toxic pharmacological pathway.

***The effect of N-acyl-glycines on thermosensitive Transient Receptor Potential (TRP) Channels:*** TRPV1, TRPV4, and TRPM3 are members of the transient receptor potential (TRP) channel family, each playing distinct roles in sensory and pain physiology. TRPV1 is a non-selective cation channel known for detecting painful heat (>43°C), acidic conditions, and chemicals such as capsaicin. It is expressed in nociceptive (pain-sensing) neurons. When activated, TRPV1 channels leads to pain signal transduction. TRPV4 is a nonselective cation channel responsive to moderate heat, osmotic changes, and mechanical stress. It plays key roles in osmoregulation, mechanosensation, and regulation of vascular function. Like TRPV1, its activation can be sensitized under inflammatory or injury conditions. TRPM3 is another polymodally-gated, nonselective cation channel sensitive to noxious heat and certain neurosteroids. It is expressed in peripheral sensory neurons and crucial for detecting high temperatures and inflammatory heat pain. Activation of TRPM3 leads to pain transmission and neuropeptide release, like TRPV1. Previously it has been shown that a mix of N-acyl-glycine species was reported to partially activate several TRP channels in the vanilloid subfamily, including the heat and capsaicin-activated TRPV1 and the osmosensitive and heat sensitive TRPV4 [22]. Here we tested the effect of NA-Gly, DHA-Gly and 19,20-EDP-Gly on agonist-induced Ca^2+^ signals in HEK293 cells transfected with TRPV1, TRPV4, and TRPM3 (another heat activated TRP channel). Application of 5 μM NA-Gly, DHA-Gly or 19,20-EDP-Gly alone did not induce any significant increase in the cytoplasmic Ca^2+^ by itself in cells expressing TRPV1, TRPV4 or TRPM3. All three N-acyl glycines however significantly potentiated Ca^2+^ signals induced by the TRPV4 agonist GSK1016790A (50 nM) in TRPV4 expressing cells (Figure 7A and 7B). GSK1016790A-induced Ca^2+^ signals developed faster, reached a higher peak, but also decayed faster in cells pretreated with N-acyl-glycines, presumably due to channel desensitization. Ca^2+^ signals induced by TRPM3 agonist 12.5 μM pregnenolone Sulfate were not affected significantly by 5 μM NA-Gly, DHA-Gly or 19,20-EDP-Gly (Fig 7C and 7D). Similarly, Ca^2+^ signals induced by 20 or 100 nM capsaicin were not altered significantly by any of the N-acyl-glycines in TRPV1 expressing cells (Figure 7E and 7F).

**Figure 7.**
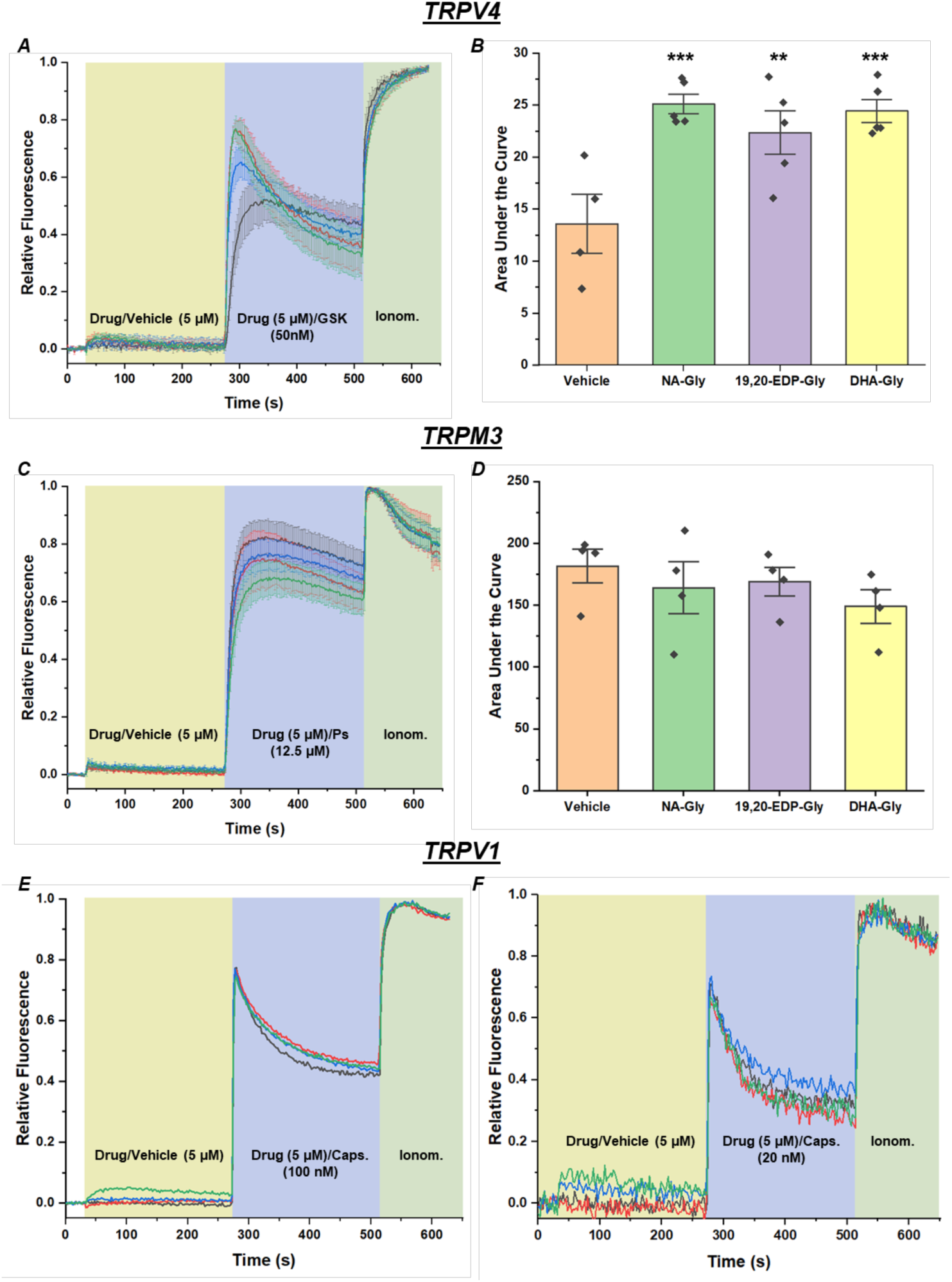
N-acyl-glycines potentiate TRPV4 activity and do not affect TRPV1 and TRPM3 activity. Cytoplasmic Ca^2+^ signals in HEK293 cells transfected with TRPV4, TRPM3, TRPV1 and GCaMP6 were measured as described in the methods section using a FlexStation 3 plate reader. **(A)** Average traces from 5 independent transfections. First 5 μM NA-Gly, DHA-Gly or 19,20-EDP-Gly, or vehicle (DMSO) was applied, followed by 50 nM GSK1016790A (TRPV4 agonist), and by 2 μM ionomycin. Error bars show SEM. **(B)** Bar graph show the area under the curve for the individual experiments in the first 40 s of the application of GSK1016790A. Statistical significance was calculated by one-way analysis of variance. Mean ± SEM and scatter plots. **(C)** First 5 μM NA-Gly, DHA-Gly or 19,20-EDP-Gly, or vehicle (DMSO) was applied, followed by 100 nM (A) or 20 nM (B) capsaicin, and by 2 μM ionomycin. **(D)** The area under the curve after the application of the N-acyl Glycines before the application of capsaicin was plotted. The differences between the different groups by one-way analysis of variance was not significant (p=0.08, overall ANOVA). **(E)** Average traces from TRPM3 transfected cells from 4 independent transfections. First 5 μM NA-Gly, DHA-Gly or 19,20-EDP-Gly, or vehicle (DMSO) was applied, followed by 12.5 uM Pregnenolone Sulfate, and by 2 μM ionomycin. **(F)** Bar graph show the area under the curve for the individual experiments after the application of Pregnenolone Sulfate. Mean ± SEM and scatter plots. The differences between the different groups by one-way analysis of variance was not significant (p=0.53 for overall ANOVA).

## DISCUSSION

Diets rich in ω-3 fatty acids are known to promote neurological health. *N*-acyl amino acids, specifically those that are docosahexaenoic acid based are gaining interest. Furthermore, the oxidative and non-oxidative metabolites, including endocannabinoids, lipidated-amino acids and lipidated-neurotransmitters can elicit beneficial health properties through various pathways or receptors. In this study, we aim to investigate the properties DHA-Gly and its epoxides with respect to their anti-inflammatory potential and ability to activate TRP channels.

We first detected and quantified the presence of both NA-Gly and DHA-Gly in porcine hippocampus and cerebellum. Although the levels at 8 pmol/g tissue is low, these are the basal levels present in healthy porcine brains without any special method of extraction and enrichment. Previously lipidated glycines have been shown to exist in brain tissues from rodents [22, 23]. While traditionally conserved traits across species are attributed to genes or sequences, the fact that these molecules are present in multiple species suggests a common physiological relevance with respect to brain function.

Like many endocannabinoids and lipidated metabolites, it is plausible that these basal levels increase following an inflammatory or pathogenic stimuli. Therefore, we measured the formation of NA-Gly and DHA-Gly in rodent microglia with or without inflammatory LPS stimuli. We observed two trends: 1) NA-Gly and DHA-Gly levels increase upon the supplementation of their own precursor molecules including glycine supplementation and/or lipid, and 2) inflammatory LPS stimuli greatly increase NA-Gly and DHA-Gly levels compared to non-inflammatory saline. The first trend is highlighted by examining non-LPS or LPS stimulated groups independently, comparing only the supplementations. For instance, within the non-LPS group, levels of NA-Gly and DHA-Gly both increase upon supplementation of glycine, parent precursor lipid, or both. The same trend applies to LPS-stimulated samples. The second trend that inflammatory stimuli further increase NA-Gly and DHA-Gly levels compared to non-LPS control is clear when examining the same supplementation group (i.e. arachidonic acid + glycine) in two different stimulation groups (i.e. non-LPS vs LPS). Our data demonstrates that an inflammatory or pathogenic insult increases the levels of lipidated-glycine molecules. These findings are corroborated by previous studies where carrageenan-induced inflammation increased levels of lipids in rodent brains [22].

In our study, we report the ability of NA-Gly and DHA-Gly to act as inhibitors of FAAH. It is well established that inhibitors of FAAH will inherently increase levels of anandamide. Although this may increase the levels of circulating anti-inflammatory molecule anandamide, it is plausible that inhibitors of FAAH are also substrates of FAAH. The hydrolysis of the amide linkage of the head group of *N*-acyl amides may present an availability of a new amide linkage forming between the head group of *N*-acyl lipids and other amino acids. Therefore, we investigated whether NA-Gly and DHA-Gly were still detected with the presence of FAAH inhibitor JZL195 in non-LPS or LPS-stimulated conditions. We showed that NA-Gly and DHA-Gly were still detected at high quantities of JZL195, suggesting that these lipidated glycine molecules are likely forming under other biosynthetic routes. Interestingly, previous studies have highlighted the significance of hydrolase enzymes having bi-functional properties including hydrolase or synthase activity [19]. Crucially, this study also reports on the cooperative enzymatic control of *N*-acyl amino acids by PM20D1, a different synthase/hydrolase found in the body. Therefore, it is plausible the ability of rodent microglia to form NA-Gly and DHA-Gly in the presence of FAAH inhibitor JZL195 is due to the presence of PM20D1.

Structurally, lipidated-neurotransmitters and lipidated-amino acids are similar to endocannabinoids. Endocannabinoids are endogenous lipid molecules that interact with and activate the cannabinoid receptors. Some of the most well-studied endocannabinoids are DHEA and AEA, both of which contain an ethanolamine moiety at the head group of docosahexaenoic acid or arachidonic acid, respectively. It was reported that the formation of epoxides of endocannabinoids such as DHEA can increase the affinity for cannabinoid receptors [26]. Therefore, we investigated whether NA-Gly and DHA-Gly were agonists of cannabinoid receptors. We report that NA-Gly and DHA-Gly are not agonists of cannabinoid receptors. Although the difference between ethanolamide-conjugated or glycine-conjugated lipids are in the head group of the carboxylic acid, it is not surprising that the agonistic properties with cannabinoid receptors would differ. While the structures are similar, these changes in the head group moiety will inherently influence the ability to access the active site or bind with the cannabinoid receptors themselves.

In this study, we employed a highly sensitive G protein nanoBRET assay to systematically evaluate the signaling interactions of two glycine-conjugated lipid mediators-docosahexaenoyl-glycine (DHA-Gly) and N-arachidonoyl-glycine (NA-Gly) across a panel of canonical and noncanonical cannabinoid receptors: CB1R, CB2R, GPR55, and GPR119. Our findings provide important insights into the receptor specificity and functional roles of these endogenous lipids, particularly in relation to their agonist and inverse agonist activities.

Using a validated BRET-based platform, we first confirmed the ability of CB1R, CB2R, GPR55, and GPR119 to engage their respective cognate G proteins upon stimulation with established synthetic agonists. This provided a robust baseline to assess the potential receptor-activating properties of DHA-Gly and NA-Gly. Surprisingly, neither lipid elicited measurable activation of any of the four receptors tested, even at saturating concentrations, and did not engage non-cognate G proteins either. These results strongly suggest that neither DHA-GLY nor NA-Gly acts as a direct agonist of canonical or atypical cannabinoid receptors, in contrast to prior assumptions or indirect observations in the literature.

However, our follow-up investigations revealed a striking functional distinction between the two lipids. While DHA-Gly remained functionally inert in our assays, NA-Gly demonstrated significant inverse agonist activity, particularly at GPR55. This was evident from the reduction in BRET signal following NA-Gly treatment of receptor-preactivated cells, with GPR55 showing the most pronounced response. The effect was receptor-specific, as evidenced by a significantly smaller reduction in BRET signal observed at the unrelated α2-adrenergic receptor. These findings were further supported by an orthogonal functional assay using an SRE-luciferase reporter sensitive to GPR55 signaling via the G12/13 pathway. NA-Gly completely abolished constitutive and agonist-induced luciferase accumulation, further reinforcing its inverse agonist profile at GPR55.

The functional specificity of NA-Gly as an inverse agonist of GPR55 is particularly compelling given the receptor’s known involvement in inflammatory and nociceptive signaling pathways. GPR55 is often referred to as an “orphan” or “atypical” cannabinoid receptor and has been implicated in several physiological and pathological processes, including pain, inflammation, and cancer cell proliferation. The ability of NA-Gly to dampen GPR55 signaling suggests that it may act as an endogenous brake on GPR55-mediated excitatory or proliferative signals, adding an important layer of regulatory complexity within the signaling system.

In contrast, the lack of significant activity of DHA-Gly at these receptors indicates that its physiological roles may be mediated through other receptor classes or signaling mechanisms. Given the known anti-inflammatory and neuroprotective roles of DHA-derived lipids, further exploration of DHA-Gly’s mechanism of action is warranted. The epoxidation of lipid molecules such as anandamide and docosahexaenoyl-ethanolamide have been previously reported to be more potent anti-inflammatory molecules [26]. Considering that NA-Gly and DHA-Gly did not appear to be cannabinoid receptor agonists, it is plausible that the epoxidation couldn’t transform these molecules into cannabinoid receptor agonists. We and others have previously demonstrated that when lipidated neurotransmitters are found in the central nervous system, there can also be a formation of epoxide metabolites. It is well established that CYP2J2 epoxygenase among other CYP450s are abundant in the brain [26, 27, 58–63]. To first identify whether NA-Gly or DHA-Gly are substrates for CYP450s, we conducted binding studies against CYP2J2 and CYP3A4. Here, we report a tight binding of NA-Gly and DHA-Gly to CYP450 epoxygenases CYP2J2 and CYP3A4, both of which are found in the brain. These binding data of lipidated-glycine to CYP450s suggests there may be formation of downstream metabolites. We report the formation of epoxide metabolites of NA-Gly and DHA-Gly as measured by untargeted mass-spectrometry of purified recombinant CYP450-mediated metabolism. To confirm these findings, we repeated these experiments with rat brain and human liver microsomes. We discovered the presence of *m/z* corresponding with the epoxide metabolites of NA-Gly and DHA-Gly which is the first study reporting the epoxide metabolite of NA-Gly and DHA-Gly.

Transient receptor potential (TRP) channels, particularly the thermo-sensitive subfamilies TRPV1, TRPV4, and TRPM3, play key roles in sensing temperature, pain, and inflammation. Prior work has reported that a mixture of N-acyl glycine species partially activated TRPV1 and TRPV4 channels, with statistical significance observed only for TRPV1. However, the study used a lipid mixture, leaving unresolved whether individual species contribute equally or selectively to channel modulation. To address this gap, we investigated the activity of three chemically distinct N-acyl glycines = N-arachidonoyl glycine (NA-Gly), docosahexaenoyl glycine (DHA-Gly), and 19,20-epoxydocosapentaenoyl glycine (19,20-EDP-Gly) on three thermo-sensitive TRP channels: TRPV1, TRPV4, and TRPM3. Interestingly, none of the individual compounds robustly activated any of these channels under our experimental conditions. While DHA-Gly and 19,20-EDP-Gly induced a modest increase in TRPV1 activity, the effects were highly variable and did not reach statistical significance. These findings suggest that the previously observed partial activation of TRPV1 may be due to combinatorial or synergistic effects of multiple N-acyl glycine species in the mixture. Notably, the previously reported efficacy of the N-acyl glycine mix on TRPV1 (∼27% of the capsaicin response) was also lower than that of other related lipids, such as N-acyl GABA and N-acyl serine.

The novel observation from our study was the significant potentiation of TRPV4 activity by all three N-acyl glycines when applied in conjunction with a submaximal dose of the potent TRPV4 agonist GSK1016790A. This effect was specific to TRPV4, as similar co-application protocols with TRPV1 and TRPM3 agonists failed to yield any enhancement or inhibition of calcium responses. This finding highlights a potential allosteric modulatory role for N-acyl glycines on TRPV4, whereby these lipids do not activate the channel directly, but enhance receptor sensitivity or efficacy in the presence of orthosteric agonists. It is noteworthy that TRPV4 is known to integrate mechanical, thermal, and osmotic stimuli, and is sensitive to changes in membrane tension and lipid composition. Thus, the observed potentiation may reflect a physiologically relevant mechanism for tuning TRPV4 responsiveness in response to endogenous lipid signaling.

The lack of direct activation but significant potentiation suggests that N-acyl glycines may function as context-dependent sensitizers rather than classical agonists. In physiological settings, such modulatory roles could enhance TRPV4 activity during inflammation or tissue injury, where both endogenous lipids and TRPV4 agonists (e.g., mechanical stretch or osmolytes) are elevated. Such a mechanism could contribute to altered pain perception or vascular tone. Finally, the specificity of this effect for TRPV4 over TRPV1 or TRPM3 underscores the need to consider channel-specific lipid interactions when exploring lipidated neurotransmitter or endocannabinoid-like molecules. While TRPV1 remains a prominent target in the context of endovanniloid signaling (e.g., anandamide and capsaicin), TRPV4 may be more susceptible to subtle allosteric modulation by endogenous N-acyl glycines. Overall, the study demonstrates that NA-Gly and DHA-Gly do not robustly activate TRPV1, TRPV4, or TRPM3 channels but can significantly potentiate TRPV4-mediated calcium signaling under submaximal agonist conditions. These findings shift the focus from direct activation to allosteric modulation and point to TRPV4 as a potentially important target of N-acyl glycine lipid signaling.

In summary, diets rich in ω-3 fatty acids support neurological health, and docosahexaenoyl-conjugated N-acyl glycines are emerging as important lipid signaling molecules. This study explored the anti-inflammatory and ion channel-modulating roles of docosahexaenoyl glycine (DHA-Gly) and N-arachidonoyl glycine (NA-Gly), as well as their epoxide metabolites. These lipids were found endogenously in porcine brain and showed increased formation in LPS-stimulated rodent microglia, suggesting a role in inflammation. NA-Gly and DHA-Gly bound to brain-expressed CYP450 enzymes (CYP2J2, CYP3A4), and their epoxide metabolites were identified in vitro and in microsomal systems, marking the first evidence of such metabolism. Despite structural similarity to endocannabinoids, both NA-Gly and DHA-Gly lacked agonist activity at CB1R, CB2R, GPR55, and GPR119, though NA-Gly exhibited inverse agonist activity at GPR55, potentially regulating inflammatory signaling. While neither NA-Gly, DHA-Gly, nor 19,20-EDP-Gly directly activated TRPV1, TRPV4, or TRPM3 channels, all three potentiated TRPV4 activity in the presence of a submaximal agonist.

Together, the findings highlight NA-Gly and DHA-Gly as bioactive lipid mediators with selective receptor and channel interactions. Their role as inverse agonists and sensitizers, rather than direct agonists, refines our understanding of lipid signaling in the brain, particularly under inflammatory conditions.

## METHODS

### Synthesis of DHA-Glycine and 19,20-EDP-Glycine

DHA-Glycine was synthesized via conjugation of docosahexaenoic acid to glycine. To DCM (6 mL), docoshexaenoic acid (1 equiv.), EDC (2 equiv.), N-hydroxyphthalamide (1.1 equiv), and DMAP (0.1 equiv.) was added. The mixture was stirred covered from light for 3 hours, and solvents were removed by rotary evaporator. Hexane (20 mL) was added, and mixture was filtered through a pad of silica, and subsequently washed with additional hexane (20 mL). The elution was concentrated by rotary evaporator and resuspended in tetrahydrofuran (12 mL). To this solution, K2CO3 (2 equiv.), H2O (12 mL), and glycine (2 equiv.) was added and mixture stirred for 3 hours. The reaction was diluted with 1M HCl (12 mL) and CH2CL2 (30 mL). The layers were separated, and aqueous phase extracted with CH2Cl2 (30 mL). The combined organic extracts were concentrated and filtered prior to purification on HPLC. 19,20-EDP-Glycine was synthesized via conjugation of 19,20-EDP to glycine. Firstly, 19,20-EDP was synthesized as previously described. Briefly, mCPBA (2 equiv.) and DHA (1 equiv.) were added to DCM (2 mL). The mixture was stirred for 1 hour at room temperature, and subsequently extracted using H2O and DCM prior to purification on HPLC. Purified 19,20-EDP was then conjugated to glycine utilizing the same conjugation method described above for the conjugation of DHA to glycine.

### Metabolism to Identify Metabolite Products

The metabolism of NA-Gly, DHA-Gly, and DHEA by various CYP450s or microsomes were performed as follows. 0.5 mL reaction mixtures containing substrate (100 μM), CYP (0.6 μM), CPR (0.6 μM), POPS:POPC lipid reconstituted system (50 μM), 0.1 M potassium phosphate buffer (pH 7.4) were pre-incubated at 37 C for 10 minutes. Reactions were initiated with 6 mM NADPH and terminated after 30 minutes with 0.5 mL of ethyl acetate. The metabolites were extraction three times with 3 mL ethyl acetate:hexane (8:2), centrifuged at 800 *xg* for 5 minutes at 4 C, and dried under steady N2 stream. Samples were then submitted for LC-MS/MS analysis as described below.

### tGFP-FAAH- and tGFP-PM20D1-mediated hydrolysis of lipidated amino acids

tGFP-FAAH was obtained from Origene (CAT#: RG210331). tGFP-FAAH is a human cDNA clone of fatty acid amide hydrolase (Accession Number: NM_001441) containing a tGFP tag. tGFP tag is a turbo GFP, which is a small but highly fluorescent GFP. tGFP-FAAH was transformed in DH5α *E. Coli*, expanded, and DNA isolated. DNA was then utilized for transfection in HEK293 cells using Calfectin (SignaGen CAT#: SL100478) according to manufacturer guidelines. Transfection was confirmed via fluorescence microscopy.

tGFP-PM20D1 was also obtained from Origene (CAT#: RG209009). tGFP-PM20D1 is a human cDNA clone of human peptidase M20 domain containing 1 (Accession Number: NM_152491). tGFP-PM20D1 was transformed in DH5α *E. Coli*, expanded, and DNA isolated. DNA was then utilized for transfection in HEK293 cells using Calfectin (SignaGen CAT#: SL100478) according to manufacturer guidelines. Transfection was confirmed via fluorescence microscopy.

To identify whether FAAH, PM20D1, or both can mediate the hydrolysis of lipidated amino acids, 10 μM NA-Gly, DHA-Gly, or Anandamide (positive control) were added. Cells with the lipidated amino acids were incubated for 24 hours at 37C and 5% CO2. The supernatants and cells were then frozen for further processing for LC-MSMS analysis.

### BV-2 Inflammation assay

BV-2 cells were grown in Dulbecco’s Modified Eagle Medium (DMEM) supplemented with 10% FBS, 1% penicillin/streptomycin containing 1mM Glutamax and 1mM sodium pyruvate in an incubator at 37C with 5% CO2. Cells were passaged and plated on 96-well plates at 5x10^4^ cells/well. Cells were pre-treated with DHEA, NA-Gly, or DHA-Gly for 4 hours, then stimulated with LPS at 25 ng/mL. After 24 hours, supernatants were collected and frozen for cytokine analysis. IL6 production was analyzed using

Invitrogen uncoated mouse IL6 ELISA kit (Thermo Fisher Scientific, CAT# 88-7064-88) according to manufacturer’s protocols. Whether these molecules were toxic were confirmed by the lactate dehydrogenase cytotoxicity assay (Cayman Chemical, CAT#: 601170) BV-2 time dependent assays were prepared in the same way as described above. Cells were plated on 10cm plates prior to stimulation. For stimulation, media on cells at 80% confluency were aspirated, and fresh media was added either with or without 10 ng/mL lipopolysaccharide. Each stimulation timepoint (with or without LPS) was set up in triplicate for each timepoint (0 hours, 0.5 hours, 1 hour, 2 hour, 4 hours, 6 hours, 12 hours, 24 hours, 48 hours). At each timepoint, the media was collected and stored at 80C for cytokine expression analysis. Cells were washed once with ice cold phosphate buffer saline (pH 7.4), and detached from the plate using 1 mL Trizol and stored at -80C for RNA isolation.

RNA isolation was performed using a Zymo kit according to manufacturer guidelines. Reverse transcription was subsequently carried out on 1 μg of RNA per samples using a high-capacity cDNA reverse transcription kit (Applied Biosciences) according to manufacturer’s guidelines. qPCR was conducted using SYBR-Green, with forward and reverse primers (0.1 μg/μL per primer). Total qPCR reaction volume was 20μL and was performed using Applied Biosystems qPCR machine with StepOne Software Version 2.3.

### LC-MS/MS Fragmentation

Ultra-performance liquid chromatography coupled to mass spectrometry (UPLC-MS) was performed using a Vanquish Horizon (ThermoFisher Scientific), fitted with a Waters Corporation ACQUITY UPLC BEH C18 column (2.1 × 100 mm, 1.7 µm particle size), coupled to a high resolution accurate mass Orbitrap Exploris 240 mass spectrometer (ThermoFisher Scientific). The chromatographic method for sample analysis involved elution with water with 0.1% formic acid (mobile phase A) and 80% isopropanol 20% acetonitrile with 0.1% formic acid (mobile phase B) using the following gradient program: 0 min 95% A; 0.5 min 95% A; 2.5 min 40% ; 8.5 min 0% A; 10.8 min 0% A; 10.9 min 95% A; 12 min 95%

A. The flow rate was set at 0.40 mL/min. The column temperature was set to 60 °C and the injection volume was 0.5 µL.

The Exploris 240 heated electrospray ionization (HESI) source was operated at a vaporizer temperature of 420 ⁰C, ion transfer tube temperature of 300 ⁰C a spray voltage of +3.5 kV, and sheath, auxiliary, and sweep gas flows of 40, 8, and 1, respectively. Full scan data, 120-1200mz, was collected in in the orbitrap at 120,000 resolution. MS/MS experiments were collected utilizing Thermo Scientific Xcalibur AcquireX deep scan data dependent acquisition (DDA). Precursor ions were isolated with a 0.8mz window, activated by normalized stepped HCD set at 15,30,50%, and analyzed in the orbitrap at 15,000 resolution.

The Exploris 240 heated electrospray ionization (HESI) source was operated at a vaporizer temperature of 420 ⁰C, ion transfer tube temperature of 300 ⁰C a spray voltage of +3.5 kV, and sheath, auxiliary, and sweep gas flows of 40, 8, and 1, respectively. Mass spec data was acquired using data-dependent acquisition (DDA) experiments. Survey full spectra were collected between 120-1200mz at 120,000 resolution and a 0.6 second cycle time. MS/MS spectra were collected by isolating precursors with a 0.8mz window, activated by normalized stepped HCD set at 15,30,50%, and analyzed in the orbitrap at 15,000 resolution. An exclusion list was generated from the top 200 background ions and dynamic exclusion was set to 1.5 seconds with a mass tolerance of 3ppm.

### LC-MS/MS Fragmentation Analysis

UPLC-MS raw data files were analyzed utilizing Thermo Scientific Xcalibur Qual Browser Software, Version 4.1.50. The filters in Table # were utilized to identify the peaks of the liquid chromatogram. The peaks on the liquid chromatogram were then individually selected to identify the molecules with corresponding target *m/z* values in the mass spectrometry chromatogram. The molecules were identified by the corresponding mass of M + adduct, where the most common adduct was H^+^.

|  | **Target $m/z$** | **Mass Range Filter for LC** | **Target Adduct** |
| --- | --- | --- | --- |
| **DHEA** | 371.8 | 370 – 375 | H<sup>+</sup> |
| **DHEA-D<sub>4</sub>** | 375.31 | 370 – 380 | H<sup>+</sup> |
| **DHEA-Epoxyde** | 387.28 | 385 – 390 | H<sup>+</sup> |
| **DHEA-D4-Epoxyde** | 391.30 | 387 – 395 | H<sup>+</sup> |
| **DHA-Gly** | 385.26 | 380 – 390 | H<sup>+</sup> |
| **Epo-DHA-Gly** | 401.26 | 400 – 405 | H<sup>+</sup> |
| **NA-Gly** | 361.26 | 300 – 367 | H<sup>+</sup> |
| **Epo-NA-Gly** | 377.26 | 375 – 385 | H<sup>+</sup> |

***Reagents for GPCR assays.*** AM-251 was purchased from Tocris (#1117) and resuspended in DMSO to a stock Concentration 10mM); GSK1292263 was purchased from MedChemExpress (#HY-12066) and resuspended to 10mM in DMSO. NanoGlo was purchased from Promega. PEI was purchased from Polysciences (23966). Clonidine was purchased from Tocris (#0690) and resuspended to 50mM stock in water.

***GPCR Cell Culture and Transfection.*** HEK293T/17 cells (RRID: CVCL_1926) were purchased from ATCC and cultured in DMEM (Gibco CAT#: 10567-014) supplemented with 10% dialyzed FBS (Biowest CAT#: S181D), non-essential amino acids (Gibco CAT#: 11140-050), penicillin 100 units/ml and streptomycin 100 µg/ml (Gibco CAT#: 15140-122), and amphotericin B 250 µg/ml (ThermoFisher CAT#: 15290-018) at 37°C and 5% CO2. Cells were routinely monitored for possible mycoplasma contamination. Two million cells were seeded in each well of 6-well plates in medium without antibiotics for 4 hours and then transfected with a 1:3 ratio of DNA plasmid (2.5 µg) and polyethylenimine (PEI; 7.5 µl) (Polysciences, 23966)). Transiently transfected cells were incubated for 16 hours, then starved in OptiMEM (Gibco CAT#: 11-058-021) 6h before being tested. For luciferase reporter experiments, cells were plated on Poly-D-Lysine (Gibco CAT#: A3890401) coated 6-well plates.

***G protein NanoBRET assay.*** Plasmid for mammalian expression of assay components were transfected as follows: 208ng of human GPCR (CB1, CB2, GPR55, or GPR119), 832ng of each Gα protein, 208ng of β1-Venus(156-239), 208ng of γ2-Venus(1-155), 13ng of masGRK3CT-Nluc, and 1031 ng of pcDNA3.1 empty vector [57]. The day after transfection, cells were serum starved, collected in 1.5mL tubes, and spun at 500*g* x 5 minutes room temperature. Cell pellet was resuspended in 250μL BRET buffer (PBS supplemented with 0.5 mM MgCl2 and 0.1% glucose) and 30μL cells were plated in white 96-well microplates (Greiner Bio-One). The nanoluc substrate furimazine (Promega CAT#N1120) was diluted 1:250 in BRET buffer and 30μL were applied to plated cells. BRET measurements were obtained using a POLARstar Omega microplate reader (BMG Labtech). All measurements were performed at room temperature. BRET signal was determined as the ratio of the light emitted by Gβ1γ2-venus (emission filter 535/30) to the light emitted by masGRK3CT-Nluc (475/30). In kinetics assays, the baseline value (basal BRET ratio) was averaged from recordings of the five seconds before agonist injection, then 60µL of 2X agonist were added and BRET signal was recorded for 5 minutes. ΔBRET was obtained by subtracting the basal BRET ratio from the maximal amplitude measured and the ΔBRET obtained on pcDNA3.1 (empty vector) transfected cells is subtracted from the receptor conditions.

For the antagonist mode, 30 µL of cells were combined with 30µL of nanoluc substrate (1:250) and read for a baseline of 2 minutes. Then, 60µL of 2X agonist were applied and recordings continued until BRET ratio reached a plateau. After that, 60µL of vehicle, 3X NA-Gly, or 3X DHA-Gly were applied and measurement continued for 100 seconds. ΔBRET was obtained as previously mentioned, and normalized to 100% of the maximal BRET ratio obtained upon agonist application.

***Luciferase Reporter Assay.*** Cells were transfected with 1.20µg of SRE-RE, 0.1µg of pRL-TK (constitutively expressed Renilla Luciferase under the control of a Thymidine Kinase promoter), and 1.20µg of pcDNA3.1 or FLAG-GPR55. The day after, cells were starved for 6h in OptiMEM and treated with 100μM AM-251 or vehicle. After incubation overnight, cells were treated for 6h with 50μM NA-Gly or vehicle. After this time, cells were collected, spun at 500g x 5 minutes and cell pellet was resuspended in 200μL of BRET buffer. 30μL of cells were plated and read for luminescence of inducible firefly luciferase by application of 30μL Firefly Luciferase buffer^2^; for luminescence of Renilla Luciferase, cells were incubated with 30μL of 1:10,000 diluted Prolume Purple (#369-250, Nanolight). The maximal luminescence value of firefly luciferase was divided by the maximal value of Renilla Luciferase, and the obtained ratios were normalized as average % of pcDNA3.1 + vehicle.

***Intracellular Ca2+ measurement to measure TRP activity.*** Intracellular Ca^2+^ measurements were performed using a Flexstation-3 96-well plate reader with rapid well injection (Molecular Devices) as described earlier [64] with some modifications. Briefly Human Embryonic Kidney 293 (HEK293) cells were purchased from American Type Culture Collection (ATCC), Manassas, VA, (catalogue number CRL-1573), RRID:CVCL_0045; cell identity was verified by STR analysis by ATCC. HEK293 cells were cultured in MEM supplemented with 10% FBS and 100 IU/ml penicillin plus 100 µg/ml streptomycin in 5% CO2 at 37°C.

HEK293 cells were transfected with GCaMP6 (0.2 μg) and mTRPM3 (0.2 μg), or TRPV4 (0.2 μg) or TRPV1 (0.2 μg) using the Effectene reagent kit (Qiagen). GCaMP6f was a kind gift from Dr. Lawrence Gaspers. After 24 hr, transfected cells were plated on poly-D-lysine coated black-wall clear-bottom 96-well plates and measurements were performed 24 hr after plating. 30 min before experiments, the MEM media was replaced with a solution containing (in mM) 137 NaCl, 5 KCl, 1 MgCl2, 2 CaCl2, 10 HEPES and 10 glucose, pH 7.4 and the plate was measured at 25°C. All drugs stock (in EtOH) and control group (EtOH) dissolved in the same solution. GCaMP6 signal was measured at excitation wavelengths 485 nm and fluorescence emission was detected at 525 nm. Sampling interval was 0.86 s and four parallel reads were performed for each condition. After the experiment they were averaged and treated as one data point for statistical analysis. For all experiments, 2 μM ionomycin was applied at the end of experiment to determine the maximum response. Each group had 3-4 individual transfection. Capsaicin and GSK1016790A (GSK101) were purchased form Sigma, ionomycin and PregS from Cayman Chemicals.

## Acknowledgements

We want to thank Yi-Chien (Tiffany) Tang for editorial assistance. The authors J.S.K., A.N.D., and A.D. are grateful for the NIH NIGMS 1R35GM152121-01 grant and Georgia Tech start-up funds for funding the research. They are also grateful to Parker H. Petit Institute for Bioengineering and Bioscience (IBB) at Georgia Tech, for providing the core facility used to acquire data. All experiment work by J.S.K was completed when they were postdoctoral fellows at Georgia Tech from 2022-2024. We are grateful to Dr. Cesare Orlandi and Dr. Luca Franchini from the Department of Pharmacology and Physiology, University of Rochester Medical Center, for the experiments with GPCR and receptor interactions. The authors (L.F. and C.O.) also thank the NIDCD/NIH grant R01DC022104 and the University Research Award (URA) to C.O. We want to thank Dr. Yevgen Yudin and Dr. Tibor Rohacs for the thermosensitive Transient Receptor Potential (TRP) Channels study. The authors T.R. and Y.Y acknowledge NIH grant R01GM093290.We thank Dr. Lucas Li, from the Duke Metabolomics Centre, for running the targeted Mass spec samples on the pig brain extractions, and also, we thank the Samuel G Moore at the Systems Mass Spectrometry Core, Georgia Institute of Technology for running our untargeted mass spec samples.

## Notes

### Competing Interest Statement

The authors have declared no competing interest.

